# Multi-modal breath measurements for biomarker discovery

**DOI:** 10.1101/2023.11.09.566302

**Authors:** Phillip J. Tomezsko, Jordan Wynn, Alla Ostrinskaya, Jessie Hendricks, Trina Vian

## Abstract

Breath contains numerous classes of compounds and biomolecules that could potentially be used as biomarkers for infectious disease as well as a range of other respiratory conditions or states. A testbed for simultaneous, multi-modal measurements was developed. Seventeen healthy subjects provided breath samples at baseline repiratory rate for particle size, lipid composition and bacterial nucleic acid composition analysis. The majority of the particles the participants exhaled at baseline were smaller than 5 μm, consistent with previous literature. The exhaled breath particulate contained lipids found in lung surfactant, indicating origin in the lung. Although bacterial DNA was not significantly higher in the exhaled breath particulate than in the environmental background, the metagenome of the breath was distinct from the environment, oral cavity and nasal passages of the participants. The low abundance of the breath microbiome limited analysis. The multi-modal breath testbed has promise for discovery of breath biomarkers and as a reference for biomarkers of different classes that are currently being used.

## Introduction

Exhaled breath contains aerosols and volatile organic compounds (VOCs). During the COVID-19 pandemic, there was renewed interest in understanding the mechanisms of transmission of respiratory infectious disease. Similarly, breath has been increasingly evaluated as a source of biomarkers that can be used for diagnostic of infectious disease as well as other respiratory conditions. Previous work on the size and concentration of breath aerosols indicates that age and current activity (i.e., exercise, singing) change the distribution of particles in breath as well as their origin in the respiratory tract. Other variables such as gender, body mass index and overall level of activity were not found to impact the size distribution of aerosols in breath. The size distribution of particles in breath containing replication-competent virus has profound impacts on efficacy of mitigation measures, which was critical knowledge that was accumulated in the COVID-19 pandemic. Large aerosols are derived mostly from the oral cavity and upper respiratory tract, so spread of viruses that replicate there can be mitigated by a variety of masks that filter larger particles. SARS-CoV-2, on the other hand, replicates mostly in the lower respiratory tract, and so highly efficacious masks or environmental controls are required for mitigation.

In addition to basic knowledge about transmission of infectious disease, the use of breath as a diagnostic has been increasing rapidly, both for respiratory infections such as COVID-19 [1][2] and tuberculosis [3] as well as chronic respiratory and non-respiratory conditions including asthma, chronic obstructive pulmonary disorder (COPD), cystic fibrosis, inebriation, cancer, neonatal jaundice, heart transplant rejection [4][5]. Measurement of volatile organic compounds (VOCs) from breath are the most developed biomarkers, but other biomarkers include exhaled breath condensate and nucleic acid collection for direct detection of pathogens [6]. Clinically evaluated tests for each category exist for COVID-19.

The lung microbiome is a source of potential biomarkers that is currently limited in accessibility. The lung microbiome has very low bioburden and is typically sampled via invasive bronchial-alveolar lavage. The low biomass of the lung is maintained through frequent physical and immunological clearance of microbes [7]. Due to rapid clearance and constant intake of new microbes, the lung microbiome is highly dynamic. However, there are patterns in the types of microbes that are enriched in the lung microbiome in the healthy state. Dysbiosis in the lung is thought to contribute to a variety of conditions including asthma, COPD, pulmonary fibrosis as well as cancer and susceptibility to other infections [7]. The lung microbiome also forms axes with the brain and gut microbiome that could be important in health. Previous studies to understand the lung microbiome from breath have been unable to identify clear patterns due to the low bioburden, however the breath microbiome was distinct from the oral and nasal microbiome in both studies.

The goal of this work was to develop a capability to combine particle measurement techniques with other collection modalities as a testbed for development of novel biomarker assays in the future. This platform can also be useful for further basic science of respiratory health and transmission of respiratory diseases. Healthy volunteers breathing at baseline for extended periods of time were included in the study. We observed a low abundance of particles > 5 μm in size, similar to other studies in healthy individuals at baseline [8]. We used LC-MS to determine that collected breath aerosols were of lung origin based on enrichment of the lung surfactant dipalmitoyl phosphatidylcholine (DPPC) as compared to eluates of cheek or nasal swabs. Additionally, we collected nucleic acid and performed qPCR and metagenomics on the bacterial populations present. We observed a distinct metagenome from nasal swabs, cheek swabs and from breath. This capability could be used to investigate case-control cohorts in order to discover and validate diagnostic biomarkers.

## Methods

The breath characteriation testbed was created to study breath analytes, including host (lipids, proteins, etc), commensal species (lung microbiome), and infectious pathogens. The testbed was designed to concurrently analyze aerosols, collect particles for chemical characterization, and collect biologic samples for qPCR, metagenomics and other biological marker analyses. The study was approved by the MIT IRB (2204000622). Participants were recruited from the Lincoln Laboratory population and were in generally good health. Each subject (n=17) participated in three identical sampling collection days, which took about 45 minutes to complete. On the day of the collection, participants were asked not to eat 2 hours preceeding the study, took a rapid antigen COVID-19 test, and filled out a screening questionnaire. If the COVID-19 test was positive, the participants were asked to reschedule. The questionnaire included questions about respiratory symptoms of illness, inhaler use, and smoking history. The only exclusionary criterion was smoking the equivalent of 1 pack/day currently or for a period of 10 years ever. Each participant was also asked to complete a rapid antigen COVID-19 test the day following the collection and report their result, along with any other change of health for the following two weeks.

### 2.1 Bulk Sampling

Participants started by rinsing their mouths with distilled water. The participants then swabbed their cheek using an IsoHelix DNA Buccal Swab (Cell Projects Ltd, Harrietsham, UK) and nose using a C&A Scientific Anterior Nares Swab (Avantor, Radnor PA, USA) twice. One of each type of swabs was placed into 1 mL DNA/RNA Shield (Zymo Research, Irvine California, USA) and the other of each into 1 mL methanol. Cheek and nasal swabs were stored at -20 °C for nucleic acid extraction.

A digital peak flow meter (Microlife, Widnau, Switzerland) was used to measure peak flow rate, and then the participant was asked to practice breathing with a peak flow rate of 65 liters per minute (LPM), the target rate for sample collection.

### 2.2 Aerosol Sampling

To analyse the aerosol characteristics and a variety of potential biomarkers, we created a multiplexed sampling testbed as shown in Figure 1. The central component of the device was a stainless-steel manifold that straightens the flow and has four isokinetic nozzles for sampling. The manifold interfaced with the participant via a twin port CPAP mask (Intersurgical, Wokingham, UK). A HEPA filter was placed on the inlet of the mask in order to filter the room air before getting to the participant and sampler. Sampling ports were attached to a Fast Mobility Particle Sizer (FMPS) model 3091(TSI, Shoreview, MN, USA), an Aerodynamic Particle Sizer (APS) model 3321 (TSI), a BioSpot-VIVAS 300-P bioaerosol sampler (Aerosol Devices, Fort Collins, CO, USA), and a PTFE membrane filter cassette (0.2 μm pore size, 37 mm diameter, 10 LPM) in line with an air blower pulling air through the sampling manifold at a rate of 30 LPM. This supplementary blower ensured areas of recirculation could not set up in the manifold and exhaust could overcome the pressure drop of the ULPA filter. The manifold was heated to 37 °C via band heaters controlled via a thermal controller (PM6C1CA, Watlow, St. Louis, MO, USA) and electrostatically-conducting sampling lines were heated with manually-controlled heater tapes. Insulation of the main manifold reduced overall power consumption. A temperature set point of 37 °C was chosen to match normal body temperature and to minimize changes in the aerosol size distribution due to condensation or evaporation.

**Figure 1.**
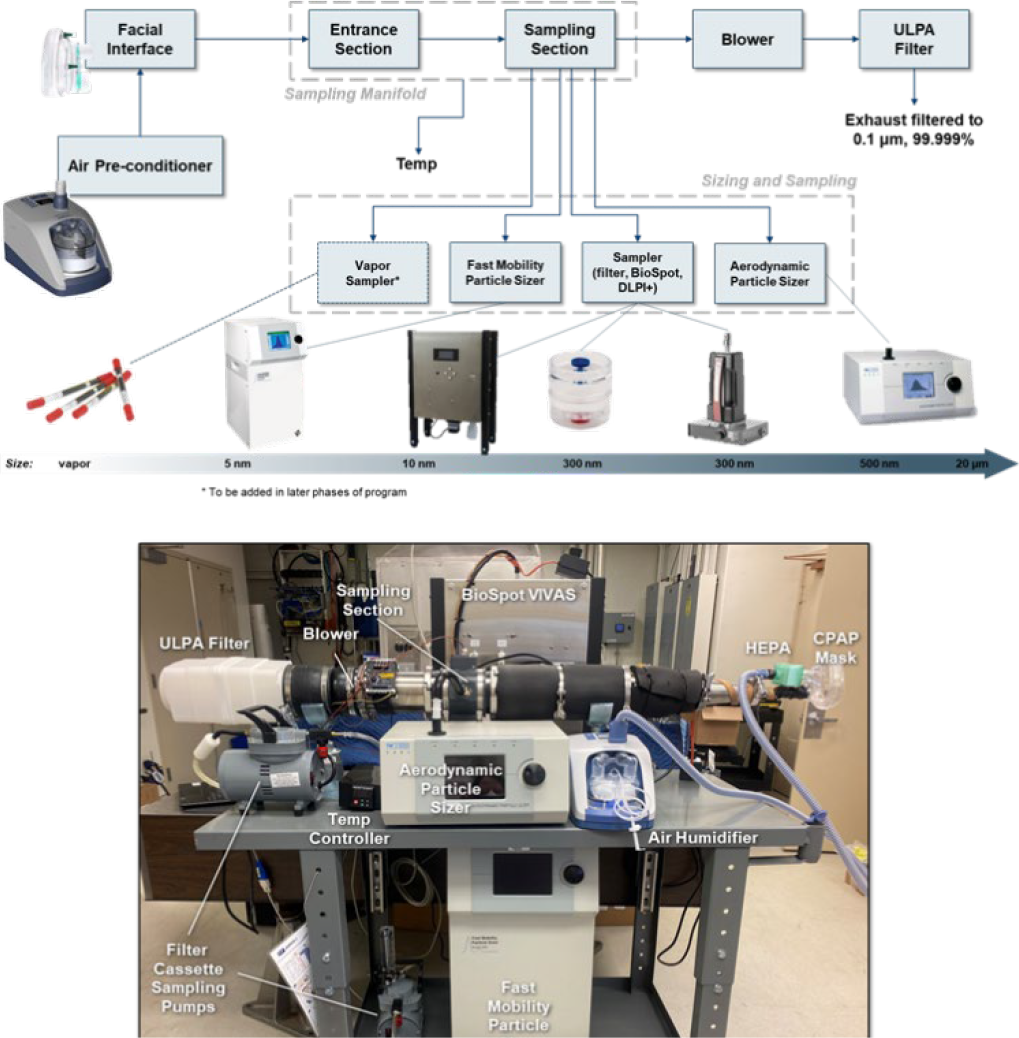
Top figure shows schematic of testbed components and collection capability. Bottom figure shows multiplexed breath sampling testbed isokinetically-sampling breath aerosol into four different instruments simultaneously.

A fresh PTFE filter cassette and BioSpot collection dish was placed on the apparatus for each sample collection. Before each session, which could include multiple participants, 5 minutes of room background was collected. After completing the rinse, swabs and peak flow measurements, participants breathed into the manifold through a fresh mask and HEPA filter for 30 minutes. Participants were free to talk during this time, though the majority of the time was spent breathing at baseline level.

#### 2.2.1 Fast Mobility Particle Sizing

The FMPS (model 3091) was connected to one of the four device ports to measure and track the distribution of sub-micron aerosol particles between 5.6 – 560 nm with a 1 Hz time resolution. The FMPS ionizes the sample and measures the electrical mobility at a sample flow rate of 10 LPM. The inlet tubing was constructed of antistatic material and heated to 37 °C.

#### 2.2.2 Aerodynamic Particle Sizing

The APS (model 3321) was connected to one of the four device ports to measure and track the distribution of 0.5 to 20 μm range particles with a 1 Hz time resolution. The sizer determines the aerosol’s diameter by calculating the time of flight of the aerosol through the test volume. The inlet tubing was constructed of antistatic material and heated to 37 °C.

#### 2.2.3 BioSpot VIVAS Sampling

The BioSpot-VIVAS 300-P bioaerosol sampler was connected to one of the four sampling ports and was used to collect and preserve aerosols from the participant, which include proteins, bacteria, metabolites, DNA/RNA, and other spores. All default temperatures and flow settings were used. The sampler works by condensing water vapor onto the aerosol, enlarging the aerodynamic diameter for efficient collection, mimicking the way the human lungs process bioaerosols. The sample was collected in a Petri-dish for further processing. Aerosols larger than 10 nm were preserved and collected through this sampling method. Samples were then collected in 2 mL DNA/RNA Shield and preserved for nucleic acid extraction.

##### 2.2.3.1 DNA extraction

ZYMOBIOMICS DNA Microprep kits (Zymo Research) were used to extract DNA according to manufacturer’s specifications. 200 μL DNA/RNA Shield solution from either the swab media or the Biospot collection was used as the input for DNA extraction. DNA was eluted in 30 μL nuclease-free water for further analysis.

##### 2.2.3.2 Quantitative PCR (qPCR)

qPCR for bacterial DNA load was performed using the Femto Bacterial DNA Quantification kit (Zymo Research). 2 μL of eluate was used as the input; samples were run in triplicate and standards were run in duplicate. The reported DNA concentration was the average of the triplicate measurements. qPCR assays were run on a Quantstudio 7 machine (Thermo Fisher Scientific, Waltham MA, USA).

##### 2.2.3.3 16S Amplicon Sequencing

The bacterial 16S rRNA V4 variable region was amplified for 5 cycles with the HotStarTaq Plus Mastermix Kit (Qiagen, Hilden, Germany). Samples were purified with Ampure XP beads (Beckman Coulter, Indianapolis, IN, USA) following manufacturer’s specifications. The libraries were sequenced at a target depth of 20,000 paired-end 150 bp reads per sample using a MiSeq machine (Illumina, San Diego, CA, USA). Reads were filtered by length <150 bp and using a maximum expected error threshold of 1.0. Reads were deduplicated, and reads with PCR errors and chimeric reads were removed. Zero-radius OTUs (zOTUs) were taxonomically classified using BLASTn against a curated database derived from NCBI [9].

#### 2.2.4 Filter Sampling

A standard, two-piece 37 mm polycarbonate personal filter cassette (PTFE filter, 0.2 um pore size) was connected to one of the four device ports and a sampling pump (10 LPM) to collect aerosol.

##### 2.2.4.1 Mass Spectrometry

PTFE filters were extracted in 2 ml of methanol, dried in a vacuum centrifuge and reconstituted into 200-μL of methanol. To account for possible loss of analytes during sample preparation, an internal standard containing isotopically-labelled d9-dipalmitoyl phosphatidylcholine (d9-DPPC; Cayman Chemical, MI USA) was added to the samples. 10 μL of sample was injected into a triple quad mass spectrometer (model API 4000, Applied Biosystems/MDS SCIEX, Framingham, MA, USA). Flow Injection Electrospray Tandem Mass Spectrometry (FIA ESI MSMS) was used for the detection of phosphatidylcholines. Quantitation was done by the external standards calibration of DPPC and PC 16:0/18:2 (Cayman Chemicals) where instrument response for the analyte was measured as a ratio between analytes and internal standard’s peak areas.

#### 2.2.8 Analysis and Statistics

Particle sizer data was processed and visualized using MATLAB (MathWorks, Natick, MA, USA) [10]. PCR, amplicon sequencing and mass spectrometry data was analysed using Excel (Microsoft, Redmond, WA, USA) [11]. Statistical analyses and visualizations were performed with R Statistical Software (R Core Team, Vienna, Austria) [12]. Principal component analysis and hierarchical clustering were performed on metagenomics data in R [12]. Dendrogram plots illustrating hierarchical clusters of samples were constructed using the dendextend R package (v1.17.1) [13].

## Results

### 3.1 Participant Demographics and Survey Scores

Seventeen healthy subjects were recruited into the study with roughly equal male/female representation. A respiratory symptom questionnaire (RSQ) was derived from [14]. The overall numeric score is the sum of reported frequency of respiratory symptoms. Scores can range from 0 to 16 (no symptoms at all to those occurring more than once daily). The mean RSQ of the population examined was 0.41 suggesting that reported respiratory symptoms were mild and infrequent overall. These demographics are summarized in Table 1.

**Table 1.** Summary of demographics for the breath sampling study.

|  |  |  |
| --- | --- | --- |
| *Number of Participants* |  | 17 |
| *Total Sampling Sessions* |  | 51 |
| *Gender (M/F)* |  | 9 / 8 |
| *COVID History (previous year)* |  | 7 |
| *Respiratory Illness History (ever)* |  | 5 |
| *Smoking History (last 10 years)* |  | 0 |
| *Age (years)* | *Mean / Median* | 40.12 / 40 |
|  | *Max / Min* | 56 / 22 |
| *Peak Flow (LPM)* | *Mean / Median* | 479 / 482 |
|  | *Max / Min* | 883 / 231 |
| *Respiratory Symptoms Questionnaire (per sample, RSQ score max = 16)* | *Mean / Median* | 0.41 / 0 |
|  | *Max / Min* | 4 / 0 |

### 3.3 Size distribution results

Breath aerosol was transported via independent sample tubes to the APS and FMPS from the sampling manifold. Total aerosol concentration for each subject session is plotted in Figure 2. APS session data from two dates during the study have been excluded because the sampling line was not heated. No correlation was found with subject’s respiratory history and exceeding the 3σ total concentration level in the APS data (Figure 2 – top). In the FMPS data, three of four subjects exceeding the 3σ level (Figure 2 – bottom) reported a history of respiratory illness.

**Figure 2.**
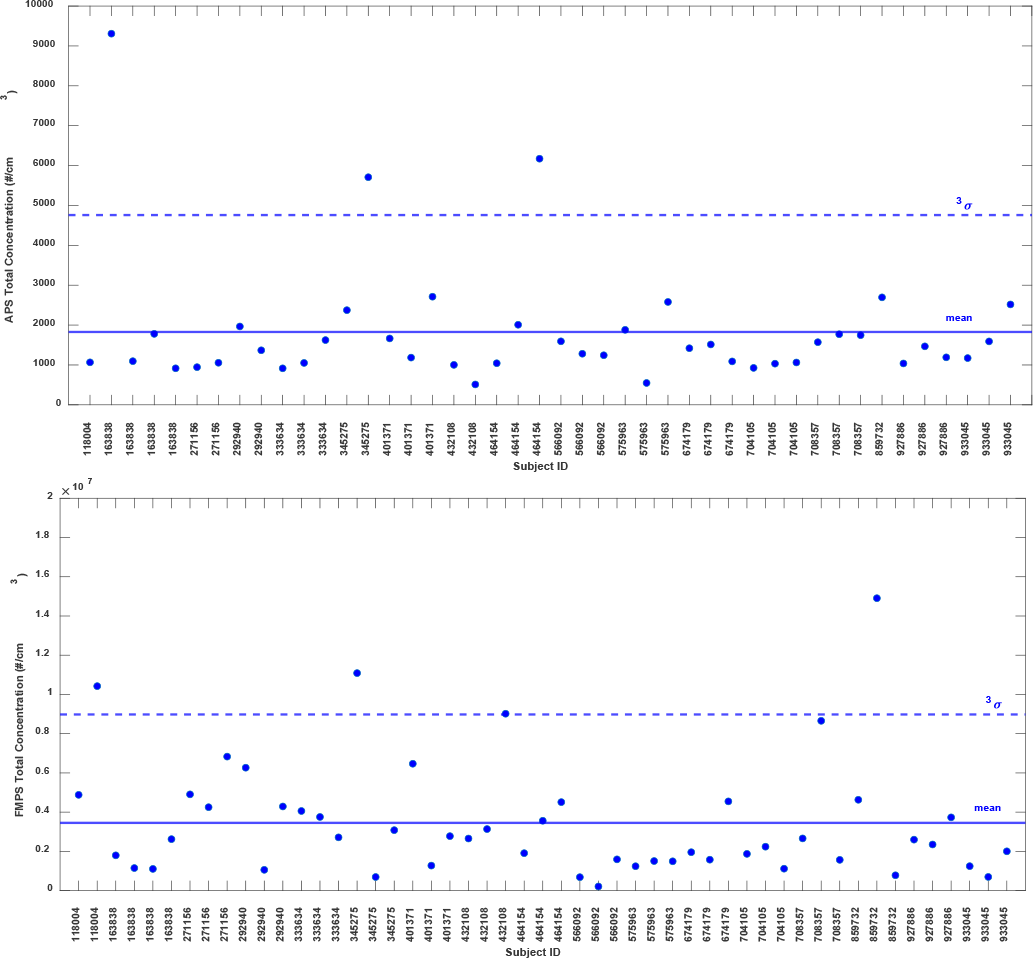
Top figure shows total aerosol concentration (#/cm^3^) recorded via APS for each session. Bottom figure shows total aerosol concentration (#/cm^3^) recorded via FMPS for each session. In both figures, mean and 3 standard deviations are denoted with horizontal lines.

The subject pool was separated into sub-groups by reported respiratory history. An additional group, “Inflammatory Event”, was created to account for those receiving vaccinations for or developing respiratory illnesses during the course of the study. Based on the guidelines of the study, only subjects testing negative for COVID and feeling well the day of sampling could contribute. Those that tested positive during the study had to wait until local quarantine periods elapsed before providing samples again. The mean aerosol size distributions for these sub-groups are shown in Figure 3. In the 0.5 to 5 μm range, little difference is noted in size distributions (Figure 3 – top). In the 5.6 to 560 nm range (Figure 3 – bottom), the location of size peaks is similar although the amplitude of the first and ratio of peaks 2 and 3 differ slightly for the sub-groups.

**Figure 3.**
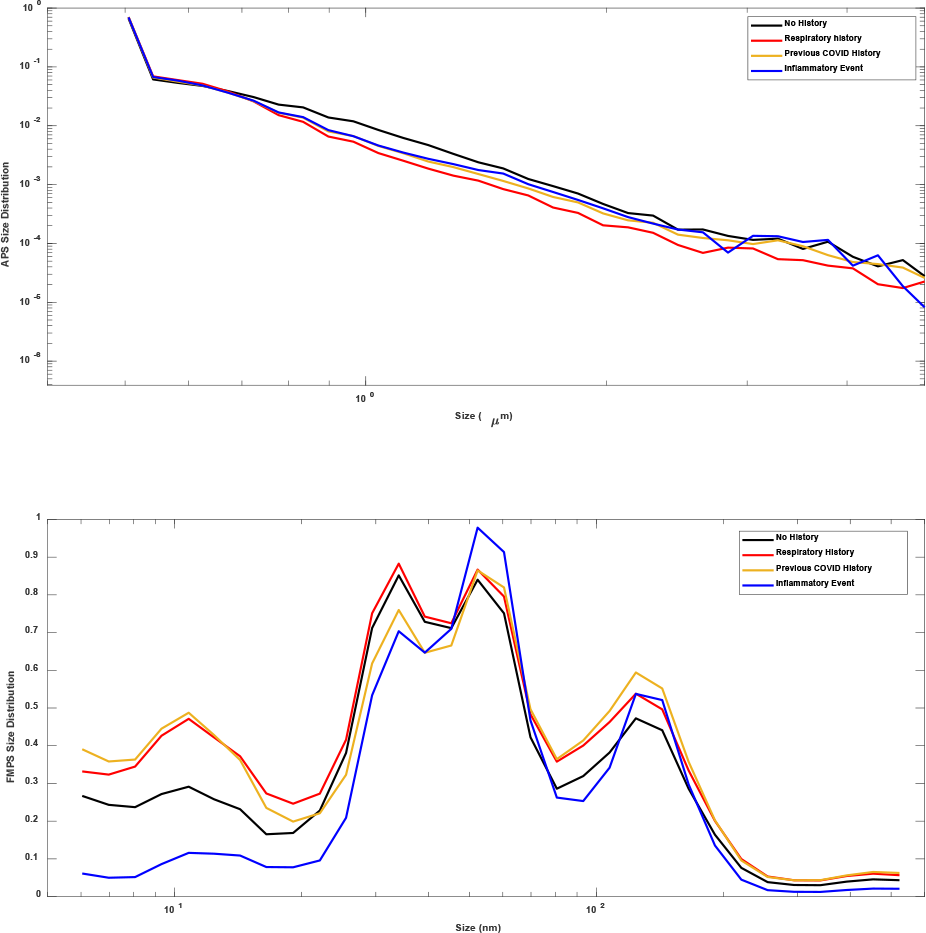
Top figure shows mean aerosol size distribution recorded via APS for each sub-group of subjects. Bottom figure shows mean aerosol size distribution recorded via FMPS for each sub-group.

One subject became symptomatic and tested positive for COVID hours after sampling. The FMPS aerosol size distribution for this subject at several timepoints during the study is plotted in Figure 4. A large shift in the proportion of aerosol at 12.4 nm is noted while the subject is pre-symptomatic. While not indicative of infectious SARS-CoV-2 aerosol, it is an indication that some shift from baseline size distribution may detectable before the onset of symptoms. More study is required to support this hypothesis. This peak remains elevated over pre-COVID levels almost two weeks later though centered at 10.8 nm.

**Figure 4.**
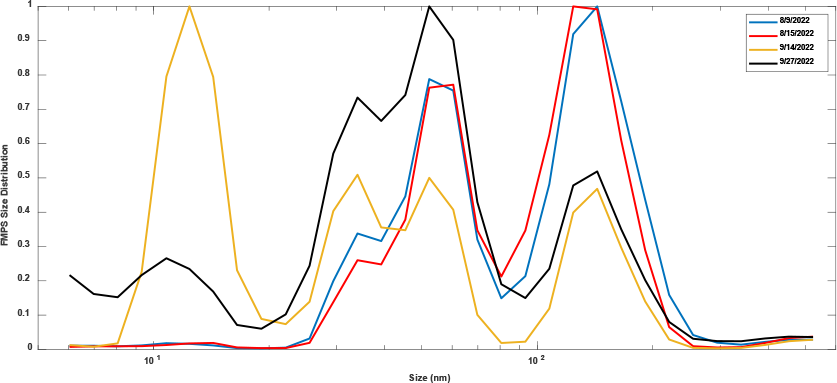
Evolution of aerosol size distribution recorded via FMPS for subject contracting COVID during study. Self-reported symptom onset occurred hours after sampling on 9/14/2022.

### 3.4 DNA results (quantity, metagenomics)

An assay for detection of bacterial DNA from low-abundance samples was selected due to the expected sparsity of DNA in breath. The limit of detection of the assay was 20 fg/μL of eluate. Detectable levels of bacterial DNA were obtained from the cheek and nasal swabs. No difference was observed in the levels of bacterial DNA in the breath BioSpot eluate samples from the room background BioSpot eluate, see Figure 5.

**Figure 5.**
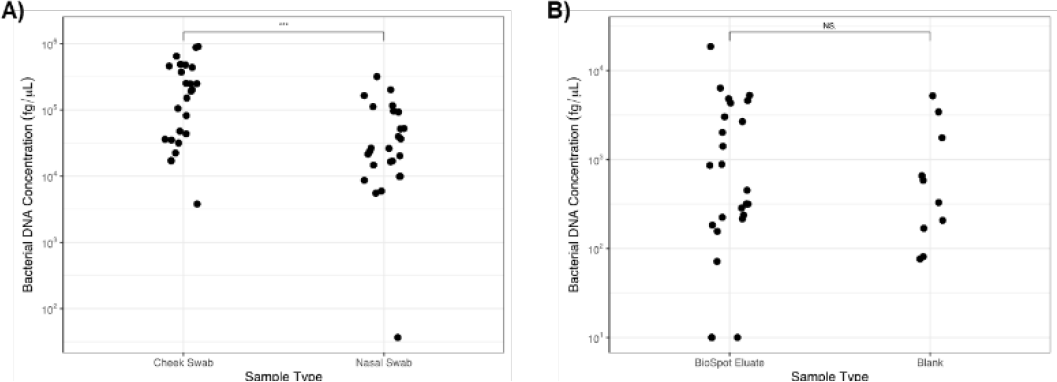
Bacterial DNA concentration. A) Bacterial DNA concentration from cheek and nasal swabs. B) Bacterial DNA concentration from BioSpot eluate during breath collection or control BioSpot eluate collected from room air (Blank). NS.-not significant, ***-p <0.001 by Wilcoxon signed ranked test

Despite the low abundance of bacterial DNA, samples from the breath BioSpot eluates were submitted for 16S amplicon sequencing. Samples from the background, nasal and cheek swabs were also sequenced for comparison. The four sample types were clustered hierarchically in order to assess relationships between the samples. We observed a clustering of a substantial fraction of the breath samples along with the background samples, indicating many of the samples were likely impacted by environmental contamination with room air, see Figure 6. A subset of breath samples clustered more closely with the nasal swab samples. However, based on the clustering with background samples and potential sources of bias including sequence batch effect, we were unable to determine if the breath samples had constituents of the lung microbiome or were combinations of room air contamination and upper respiratory microbiome. This finding matches with other attempts to identify the lung microbiome from breath that were hampered by the lung abundance of DNA and potential sources of bias or contamination [15][16].

**Figure 6.**
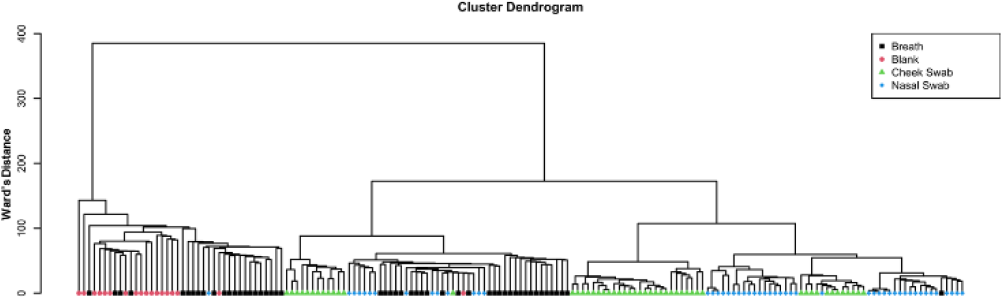
Cluster dendrogram illustrating hierarchical clustering of samples based on metagenomics measurements. Samples are categorized by type: Breath (black square), Blank (red circle), Cheek Swab (green triangle), Nasal Swab (blue diamond). The length of the branch connecting two samples or two clusters of samples indicates their level of similarity.

### 3.5 MS results

In order to verify collected breath aerosols originated in the lung, an analysis of lipid content was performed. Phosphatidylcholine (PC) 18:1/16:0, a common membrane lipid, and dipalmitoyl phosphatidylcholine (DPPC), a lung surfactant, were quantified from participant swabs and filter cassettes via mass spectrometry utilizing the procedure described by Ullah et al. [17]. No detectable lipid was measured from blank samples or unused filters and swabs. Additionally, no lipid was detected from filters which sampled room air.

A higher ratio of DPPC/PC 16:0/18:1 was observed in the filter eluate from the breath collections compared to both the cheek and nasal swabs, see Figure 7. The average DPPC/PC 16:0/18:1 from the breath filters was 1.18, which was lower than the reported average value 1.98 [9], but there was no significant difference between the ratios in this study and the previously published study (p=0.13). The findings were consistent with presence of lung surfactant in the breath collection but not in the cheek or nasal swab samples.

**Figure 7.**
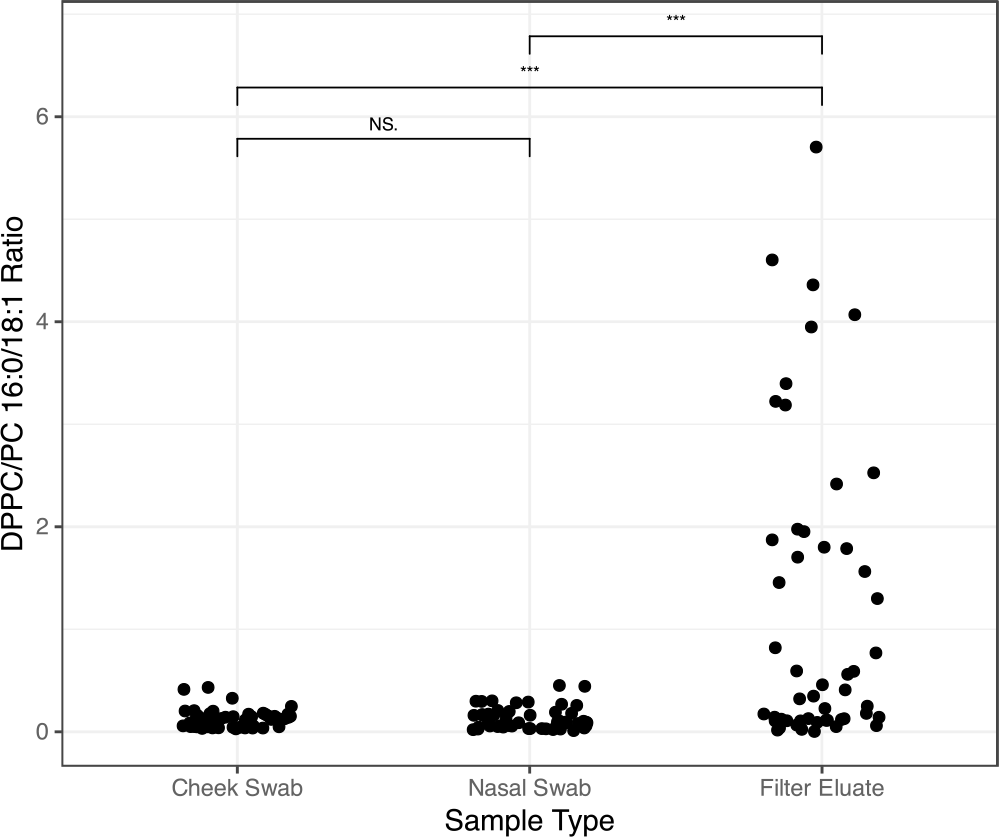
DPPC/PC 16:0/18:1 ratio determined by LC-MS. Measurements were taken from cheek swabs, nasal swabs or filters from the breath collection that were eluted in methanol. NS.-not significant, ***-p <0.001 by Wilcoxon signed ranked test

## Discussion

Other research has indicated that the size of the particle is dependent on its anatomical origin [8] thus the capability of the breath testbed to fractionate based on the particle size makes it a very valuable tool for breath research. In addition, change in the DPPC/PC 16:0/18:1 ratio with size of the particle, would support the previous findings. Nucleic acid analysis of the fractionated particles could also be useful in determining the replication site of a pathogen within the respiratory tract.

One of the hypotheses tested in this work was that the lung microbiome contributes to the breath microbiome, so breath metagenomic samples should be distinct from metagenomic samples from cheek and nasal swabs. Previous work had shown the breath microbiome is distinct from the lung microbiome, but did not show if they were more or less related than other respiratory pathway samples [15]. The current work is largely confirmatory of two other attempts to characterize the lung microbiome from breath samples, although in both cases exhaled breath condensate was sampled. In both cases, there were differences in the breath metagenome compared to samples from other collection types, but the low abundance of DNA in breath led to inconsistent results in which contamination from various sources (prep kit, sequencing batch, environment) confounded analysis [15][16]. Our metagenomic analysis showed distinct hierarchical clustering of breath samples from the background, cheek and nasal swabs. However, we could not rule out the impact of contamination and bias in our samples, as the breath samples clustered into distinct groups at least partially driven by sequencing batch effect. Our analysis does not rule out the lung microbiome contributing to the metagenome of the breath; however, a more controlled environment and higher yield collection is required.

The breath testbed can be used in the future for research into infectious disease and respiratory health in clinical populations. One potential application is for discovery and validation of biomarkers. For example, in the development of a VOC based diagnostic, the VOC profile could be compared to other sample types directly.

## Conclusions

The breath testbed is a capability that could be used for in depth, multi-modal sampling of breath for many purposes. Most importantly, it could be used to discover and/or validate biomarkers from breath. Sampling from healthy individuals at baseline confirmed that the majority of emitted particles are smaller than 5 μm. We also confirmed using MS analysis of the ratio of DPPC/PC16:0/18:1 that the particles in breath at baseline originate in the lung. We did not detect significantly more bacterial DNA in breath compared to background. Interpretation of the metagenomic sequencing of the breath was limited due the low abundance. Breath is a valuable source of distinct biomarkers that have great promise in non-invasive monitoring.

## Data Access

Data not available in the supplementary files are available upon request to the corresponding authors.

## Ethics Statements

This study was approved by the MIT IRB, the Committee on the Use of Humans as Experimental Subjects (2204000622) and the U.S. Air Force Human Research Protection Office.

All participants gave written informed consent to participate in the study.

## Supplemental Data

Refer to S1.xlsx

**DISTRIBUTION STATEMENT A. Approved for public release. Distribution is unlimited**.

**This material is based upon work supported by the Department of the Air Force under Air Force Contract No. FA8702-15-D-0001. Any opinions, findings, conclusions or recommendations expressed in this material are those of the author(s) and do not necessarily reflect the views of the Department of the Air Force**.

